# Distinct roles of the Na^+^ binding sites in the allosteric coupling mechanism of the glutamate transporter homolog, Glt_Ph_

**DOI:** 10.1101/2021.11.29.470416

**Authors:** Erika Riederer, Pierre Moenne-Loccoz, Francis I. Valiyaveetil

**Author notes:** Francis I. Valiyaveetil **Email:**. **Author Contributions:** E.R. and F.I.V. designed research; E.R. performed research; P.M-L. and F.I.V. supervised research; E.R. and F.I.V. wrote the paper.

## Abstract

Glutamate transporters carry out the concentrative uptake of glutamate by harnessing the ionic gradients present across cellular membranes. A central step in the transport mechanism is the coupled binding of Na^+^ and substrate. The sodium coupled Asp transporter, Glt_Ph_ is an archaeal homolog of glutamate transporters that has been extensively used to probe the transport mechanism. Previous studies have shown that hairpin-2 (HP2) functions as the extracellular gate for the aspartate binding site and plays a key role in the coupled binding of sodium and aspartate to Glt_Ph_. The binding sites for three Na^+^ ions (Na1-3) have been identified in Glt_Ph_ but the specific roles of the individual Na^+^ sites in the binding process has not been elucidated. In this study, we developed assays to probe Na^+^ binding to the Na1 and Na3 sites and to monitor the conformational switch in the NMDGT motif. We used these assays along with a fluorescence assay to monitor HP2 movement and EPR spectroscopy to show that Na^+^ binding to the Na3 site is required for the NMDGT conformational switch while Na^+^ binding to the Na1 site is responsible for the partial opening of HP2. Complete opening of HP2 requires the conformational switch of the NMDGT motif and therefore Na^+^ binding to both the Na1 and the Na3 sites. Based on our studies we also propose an alternate pathway for the coupled binding of Na^+^ and Asp.

## Introduction

Transport of molecules across cellular membranes is essential for life. Na^+^ coupled transporters use the existing Na^+^ gradient to carry out transport across cell membranes of a variety of substrates such as amino acids, sugars, nucleosides, neurotransmitters and vitamins.(1, 2) High resolution structural studies have revealed the molecular architecture of a number of these transporters. However, the mechanism by which the transport of the substrate is coupled to the Na^+^ gradient is not well-understood. Here, we investigate this mechanism of Na^+^ substrate coupling in the context of glutamate transporters.

Glutamate transporters, also known as excitatory amino acid transporters or EAATs are responsible for the uptake of glutamate following release into the synaptic space during synaptic transmission.(3-5) Glutamate transporters harness energy from pre-existing gradients of Na^+^, K^+^ and H^+^ across the neuronal membranes to drive the rapid uptake of glutamate.(6, 7) Rapid removal of glutamate from the synaptic space is required for efficient synaptic transmission and for preventing excitotoxicity.(3) Studies on the archaeal homologs Glt_Ph_ and the closely related Glt_Tk_ have greatly contributed to our present knowledge of the structure, dynamics and functional mechanisms of glutamate transporters.(8, 9) Glt_Ph_ and Glt_Tk_ are archaeal sodium coupled Asp symporters in which the transport of Asp is coupled to the movement of three Na^+^ ions.(10) Glt_Ph_ is a homotrimer in which each subunit consists of two domains: a trimerization domain and a transport domain (Figure 1A, B).(9, 11) The binding sites for Asp and the three Na^+^ ions (labelled Na1, 2 and 3) are contained within the transport domain (Figure 1C).(12, 13) Studies on Glt_Ph_ have shown that the transport mechanism involves the elevator like movement of the transport domain to ferry the bound Asp and Na^+^ ions across the membrane.(11, 14-17) An essential feature of the transport mechanism in Glt_Ph_ is that the binding of Na^+^ and Asp are tightly coupled.(18, 19) Structures of Glt_Ph_ and Glt_Tk_ show no direct contact between the Na^+^ ions and the bound Asp, which indicates that the mechanism of coupling involves allosteric changes in the transporter (Figure 1C). The transport domain consists of two hairpin loop regions referred to as HP1 and HP2.(12) HP2 acts as a gate to controls access to the substrate binding site when the transporter is in the outward facing state. Structural studies show that HP2 can be closed or open. The structure of Glt_Tk_ in the Apo state shows that HP2 is closed, a structure of Glt_Ph_ with only Na^+^ bound shows an open HP2 while the structures of Glt_Ph_ and Glt_Tk_ with Asp and Na^+^ bound show a closed HP2 (Figure 1C).(12, 13, 20) The structural studies and biophysical investigations suggest the following sequence of events in the coupled binding of Na^+^ and Asp (Figure 1D).(13, 19, 21) Central to the coupling mechanism is the movement of HP2. In the Apo state, HP2 is closed thereby preventing access of Asp to the binding site. The binding of Na^+^ ions to the Na1 and the Na3 sites results in opening HP2. The opening of HP2 is accompanied by a remodeling of the binding site for high affinity coordination of Asp. The binding of Asp to the transport domain is followed by the closure of HP2 and the binding of the third Na^+^ ion to the Na2 site. The transport domain with the bound Asp and three Na^+^ ions, undergoes an elevator like movement to an inward facing state from which Asp and Na^+^ ions are released into the cell. A variation of this scheme, based on computational studies, suggests that HP2 is dynamic in the Apo state and that the binding of the first two Na^+^ ions stabilizes the HP2 in the open state.(20) There are important aspects of the binding mechanism of Na^+^ and Asp to Glt_Ph_ that are not well understood. A key uncertainty is the mechanism by which Na^+^ binding to the Na1 and the Na3 sites is linked to HP2 movement.

**Figure 1:**
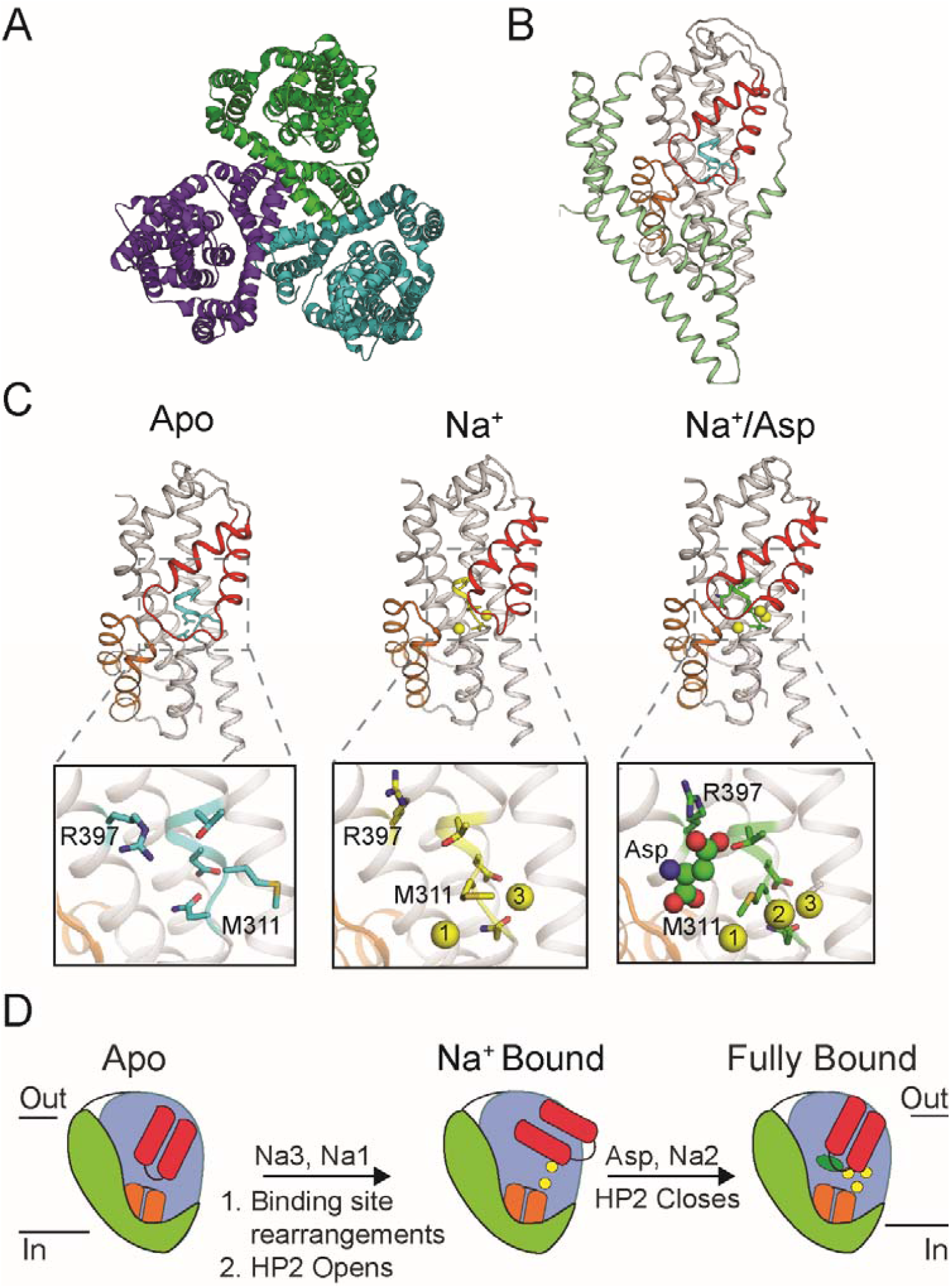
Coupled binding of Na^+^ and Asp to Glt_Ph_. **A**) Top view of the Glt_Ph_ trimer (pdb: 2nwx) with each protomer shown in a different color. **B**) A single Glt_Ph_ protomer is shown with the trimerization domain in green and the transport domain in grey. The hairpin regions are colored orange (HP1) and red (HP2) while the NMDGT motif is colored cyan. **C**) Structures of the transport domain in the Apo (pdb: 5dwy), Na^+^ bound (pdb: 7ahk) and the Na^+^/Asp bound states (pdb: 5e9s). Apo and the Na^+^/Asp structures are of Glt_Tk_ and are shown with Glt_Ph_ numbering. Insets are a close-up of the Na^+^/Asp binding sites. HP2 has been removed for clarity and key residues are shown in stick representation. **D**) Cartoon representation of the scheme for the coupled binding of Na^+^ ions (yellow spheres) and Asp (green oval).

Here, we assess the roles of the Na1 and the Na3 sites in Glt_Ph_. We develop fluorescence-based assays to track Na^+^ binding and to monitor a conformational change in the highly conserved NMDGT motif. We perturb the Na1 and the Na3 sites and use these assays along with a previously developed assay to track HP2 movement to define the specific functional roles of Na^+^ coordination at Na1 and the Na3 sites. We find that Na^+^ binding to the Na1 site results in a partial opening of HP2 while Na^+^ binding to the Na3 site results in a conformational switch in the NMDGT motif. Na^+^ binding to both the Na1 and the Na3 sites is required for a full opening of HP2 and for high affinity binding of Asp to Glt_Ph_.

## Results

### Tyr fluorescence of Glt_Ph_ reports on Na^+^ binding to the Na1 and the Na3 sites

We initially identified an assay for monitoring Na^+^ binding to the Na1 and the Na3 sites in Glt_Ph_. It has been reported that Na^+^ binding to Glt_Ph_ can be monitored by changes in protein fluorescence.(22) Glt_Ph_ does not contain any Trp residues but contains 18 Tyr residues that are distributed throughout the protein (Figure 2A).(9) The Tyr fluorescence of Glt_Ph_ shows a ∼10% increase in emission intensity with Na^+^(Figure 2B, C). This effect is selective for Na^+^ as no change in fluorescence is observed with K^+^ (Figure 2C). The change in fluorescence is also specific for Na^+^ binding as addition of Asp following Na^+^ (which causes closure of HP2) does not result in a further change in fluorescence (Figure 2C). Of the 18 Tyr residues in Glt_Ph_, three residues, Y88, Y89 and Y247 are in the vicinity (< 7 Å) of the Na1 and the Na3 sites (Figure 2A). To probe whether the fluorescence changes in Glt_Ph_ reflect Na^+^ binding to the Na1 and the Na3 sites, we substituted these Tyr residues with Phe. As a control, we also generated a Phe substitution at Y317 that was distant from the Na^+^ binding sites. We observed that the Y88F and the Y89F substitutions reduced the fluorescence change with Na^+^ while no fluorescence changes with Na^+^ were observed for a Y88F/Y89F double mutant (Figure 2D). We could not evaluate the effect of Y247 on the fluorescence change due to poor biochemistry of the Y247F mutant while the Y317F substitution, which is distant from the Na1 and the Na3 sites, did not affect the fluorescence changes observed with Na^+^ (Figure 2D). These results indicate that the changes in Tyr fluorescence of Glt_Ph_ reflect Na^+^ binding to the Na1 and the Na3 site and that changes in Tyr fluorescence provides a facile assay to monitor Na^+^ binding to these sites.

**Figure 2:**
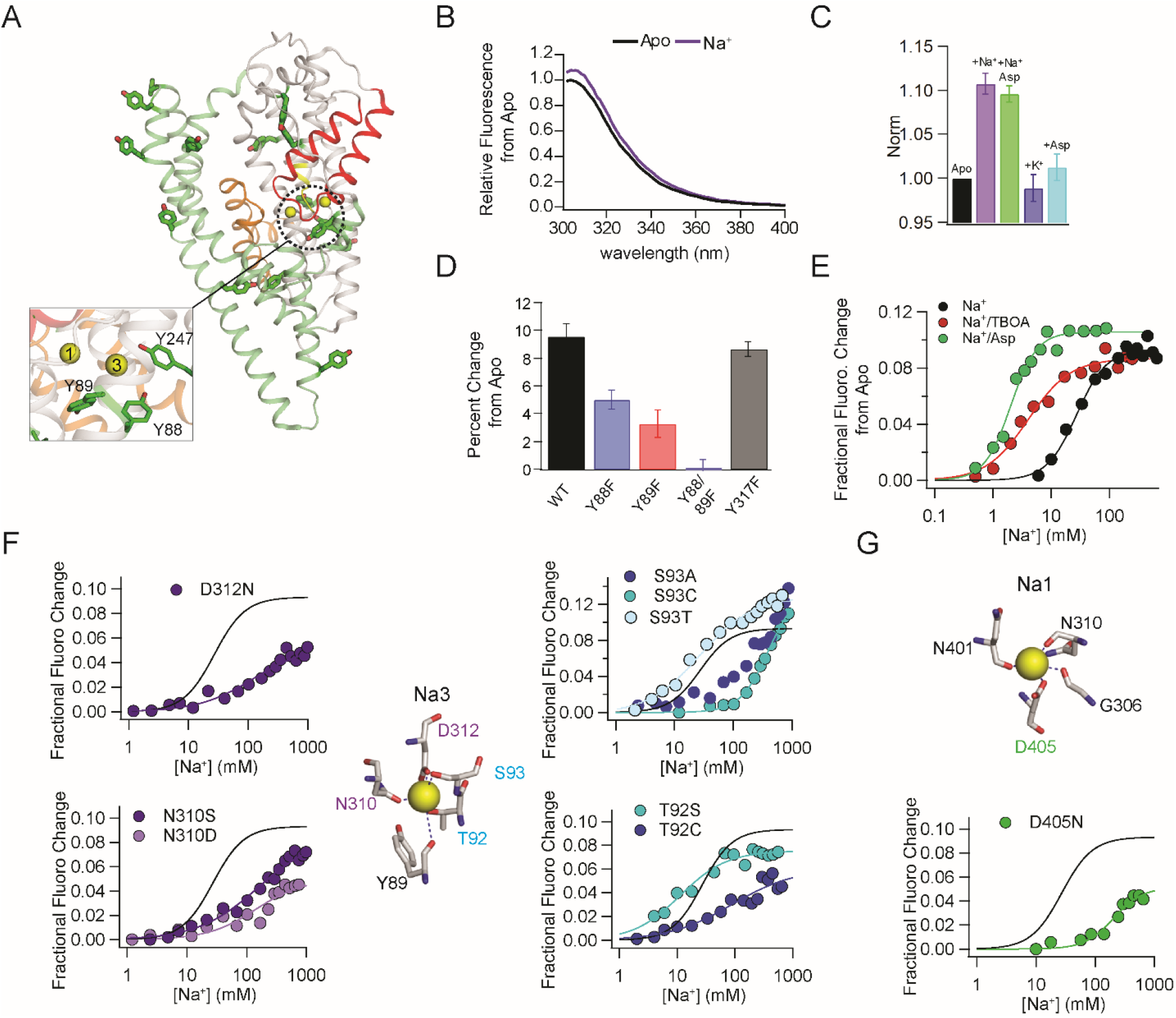
Using Tyr fluorescence to monitor Na^+^ binding to Glt_Ph_. **A**) A Glt_Ph_ protomer with Tyr residues shown in stick representation. Inset shows a close-up of the Na1/3 sites and the Tyr residues in close proximity to these sites. **B**) Fluorescence emission spectra of wild type Glt_Ph_ on excitation at 289 nm in the Apo state and after the addition of 300 mM NaCl. **C**) Fluorescence emission at 308 nm (excitation = 289 nm) for wild type Glt_Ph_ with the addition of 300 mM NaCl (purple), 300 mM NaCl/100 µM Asp (green), 300 mM KCl (blue), 100 µM Asp (teal). The fluorescence values are normalized to the Apo protein. **D**) Change in fluorescence from the Apo state for the wild type-Glt_Ph_ and Phe mutants after the addition of 300 mM NaCl/100 µM Asp. Error bars in C and D are S.E.M, n ≥ 3. **E**) Na^+^ titration of wild type Glt_Ph_. The normalized fluorescence change at 308 nm on addition of Na^+^ (black) is plotted. Sample Na^+^ titrations in the presence of 10 µM TBOA (red) and 100 µM Asp (green) are also shown. The solid line represents a fit to the Hill equation used to determine the 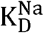. (Na^+^, 45 mM; Na^+^ with TBOA, 4 mM; Na^+^ with Asp, 1.2 mM). **F**) Close up view of the Na3 site and sample Na^+^ titrations of the various Na3 site mutants. **G**) Close up view of the Na1 site and Na^+^ titration of the D405N mutant is shown. In **F** and **G**, Na^+^ titration for the wild type Glt_Ph_ is indicted by the solid black line.

Using the Tyr fluorescence assay, we measured a 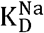 of 45 mM for Na^+^ binding to Glt_Ph_ (Figure 2E). A 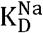 of 90 mM has been reported using a dye-based assay (18). Na^+^ binding to Glt_Ph_ has also been evaluated by the HP2 movement assay to give a value of 178 mM (19) and 120 mM by using a F273W substitution (23). The affinity of Na^+^ was increased to 1.2 mM in the presence of 100 µM Asp, as expected for a coupled transporter (Figure 2E).

To perturb the Na1 and the Na3 sites, we substituted the amino acid side chains that coordinate Na^+^ (Figure 2F, G). At the Na3 site, Na^+^ is coordinated by the side chains of T92, S93, N310 and D312. We introduced both conservative and non-conservative substitutions at these positions and evaluated the effect on Na^+^ binding. For the conservative substitutions T92S and S93T, we observed an improvement in Na^+^ binding affinity (Figure 2F, Table S1). The non-conservative substitutions showed two distinct effects. Substitutions at T92, N310 and D312 showed a dramatic shift in 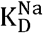 and the maximal fluorescence response observed was roughly half the value observed for the wild type protein (Figure 2F, Table S1). For the S93A/C mutants, we also observed a shift in the Na^+^ binding affinity but the maximal fluorescence response was similar to the wild type. These data support the T92C, D312N, and N310D/S substitutions abrogating Na^+^ binding to the Na3 site while the S93 substitutions decreases the Na^+^ binding affinity at the Na3 site.

At the Na1 site, D405 is the only sidechain involved in Na^+^ coordination (Figure 2G). It has been previously demonstrated using structural approaches and Asp binding assays that the D405N substitution perturbs Na^+^ binding to the Na1 site.(12) We tested Na^+^ binding to the D405N substitution using the Tyr assay. We observed a decrease in the maximal response and a shift in the 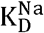 to 214 mM confirming the perturbation of Na^+^ binding to the Na1 site in the D405N mutant (Figure 2G, Table S1). We also used the dye-based assay to monitor Na^+^ binding in these Na1 and Na3 site mutants (Figure S1). The dye-based assay also reported perturbed Na^+^ binding in the Na^+^ site mutants consistent with the data obtained by the Tyr fluorescence assay. These experiments provide us with Glt_Ph_ mutants in which Na^+^ binding to the Na1 or the Na3 sites is perturbed.

### Perturbation of the Na3 site affects HP2 movement

Next, we investigated the effect of perturbing the Na^+^ sites on HP2 movement. We have previously developed a fluorescence assay to monitor the movement of HP2 (Figure 3A).(19) This assay involves the incorporation of two probes into Glt_Ph_, a Trp in HP2 and a Phe_CN_ in HP1. We substituted V355 in HP2 with Trp and used nonsense suppression approaches to substitute S279 in HP1 with cyanophenylalanine (Phe_CN_). In the Apo state, HP2 is (predominantly) closed and the probes are close together. In this state Phe_CN_ quenches the Trp fluorescence due to the proximity of the two probes (Figure 3B).(24) The binding of Na^+^ opens HP2, which increases the separation between the probes and increases Trp fluorescence due to reduced quenching of Trp by Phe_CN_. Binding of Asp closes HP2, which decreases the separation between the probes and results in a decrease in fluorescence (Figure 3B). Monitoring the changes in fluorescence therefore provides us with a means to evaluate the opening of HP2 with Na^+^ and the closure of HP2 with Na^+^/Asp

**Figure 3:**
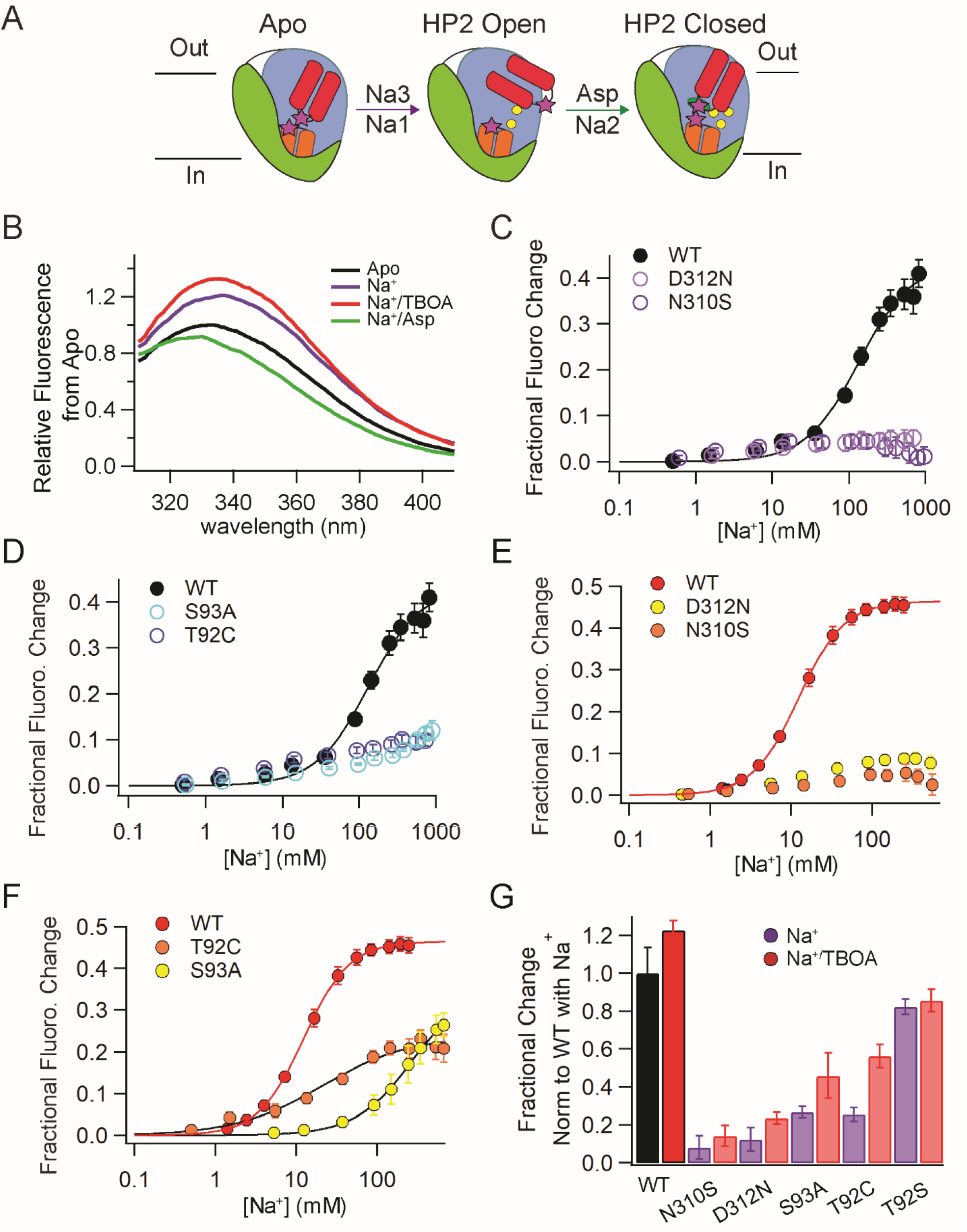
Effect of Na3 site mutations on HP2 movement. **A**) Cartoon scheme for HP2 (red) movements on binding Na^+^ and Asp. The movement is monitored by the probes (magenta stars), Phe_CN_ at 279 on HP1 (orange) and Trp355 on HP2. **B**) Emission spectra of WT Glt_Ph_ (Phe_CN_+W) on excitation at 295 nm in the following conditions: Apo, 200 mM NaCl, 200 mM NaCl/10 µM TBOA and 200 mM NaCl/100 µM Asp. **C, D**) HP2 opening by Na^+^ in the Na3 site mutants. Steady-state titration of the WT and the Na3 site mutants in the (Phe_CN_+W) background is shown. **E, F**) HP2 opening by Na^+^ in the Na3 site mutants and WT GltPh in the presence of 10 µM TBOA. Solid lines represent a fit to the Hill equation used to determine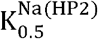. (WT: 132 ± 20 mM, WT with TBOA: 12.3 ± 0.2 mM, T92C with TBOA: 21 ± 5.3 mM and S93A with TBOA: 272 ± 33 mM). **G**) The maximal fluorescence change observed for various Na3 site mutants in the (Phe_CN_+W) background on titration with Na^+^ and with Na^+^ in the presence of 10 µM TBOA. The fluorescence changes observed are normalized to the fluorescence change observed for WT-Glt_Ph_ (Phe_CN_+W) on titration with Na^+^. Errors and error bars correspond to S.E.M for n ≥ 3.

We tested the effect of the Na3 site mutants on HP2 movement. For the N310S and the D312N mutants, we observed very small (3-5%) fluorescence changes with Na^+^ in the HP2 assay (Figure 3C, G). The lack of a change in fluorescence in the HP2 assay can be attributed to either HP2 not opening or HP2 not being propped open on the binding of Na^+^. To distinguish between these possibilities, we tested the effect of TBOA on HP2 opening. TBOA binds to Glt_Ph_ similarly to Asp but keeps HP2 propped open due to the presence of the benzyloxy group in the side chain.(12, 25) Addition of Na^+^/TBOA to the wild type Glt_Ph_ gives an additional 9% increase over the fluorescence changes seen with Na^+^ alone (Figure 3B, G). Addition of Na^+^/TBOA to the N310S or the D312N mutants showed fluorescence changes that were only slightly higher than the changes observed with Na^+^ alone, which indicates that HP2 opening is severely perturbed in these mutants (Figure 3D, G). For S93A and T92C, we observed fluorescence changes with Na^+^ and with Na^+^/TBOA indicating HP2 movement (Figure 3E, F). The changes observed for these mutants were however smaller than the changes observed in the wild type control indicating impaired HP2 opening in these mutants (Figure 3G). These results indicate that Na^+^ binding to the Na3 site is required for HP2 opening and that the magnitude of the effect depends on the specific side chain that is perturbed. We see a severe effect for substitutions in the N310 and D312 residues that are in the NMDGT motif compared to substitutions at T92 and S93 in the TM3 helix.

### A conformational switch in the NMDGT motif couples Na^+^ binding to HP2 movement

A highly conserved sequence in EAATs is the NMDGT motif.(4) This motif forms the un-wound region present in the middle of TM7 and contributes to the Na1 and Na3 sites (Figure 4A).(12) The NMDGT region shows a different conformation in Apo-Glt_Ph_ compared to Na^+^- or Na^+^/Asp-Glt_Ph_ and therefore we speculated that the NMDGT motif couples Na^+^ binding to the opening of HP2.(13, 20, 21, 26) To investigate, we developed an assay to monitor the conformational switch in the NMDGT motif. We identified that residue L99 in the vicinity of the NMDGT motif undergoes a change in environment with the conformational switch in this motif. We substituted L99 with Trp to serve as a fluorescence reporter for the NMDGT conformation switch. The L99W-Glt_Ph_ showed similar biochemistry to the wild type transporter and was functional in Asp uptake (Figure S2). We observed that the fluorescence emission of L99W-Glt_Ph_ was sensitive to Na^+^ and showed a 21% decrease in fluorescence intensity and a blue shift in the emission maximum on the addition of Na^+^ (Figure 4B, C). The blue shift in the fluorescence intensity indicates a shift to an environment with a lower polarity. There were no further changes in fluorescence on the addition of Asp, which is anticipated as there are no changes observed in the NMDGT region between the Na^+^ bound and the Na^+^/Asp bound states (Figure 4B, C). The decrease in fluorescence for L99W-Glt_Ph_ on the addition of Na^+^ was fit with a Hill equation to give a 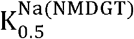 of 172 mM, which decreased to 2 mM in the presence of 100 µM Asp (Figure 4C, Table S2).

**Figure 4:**
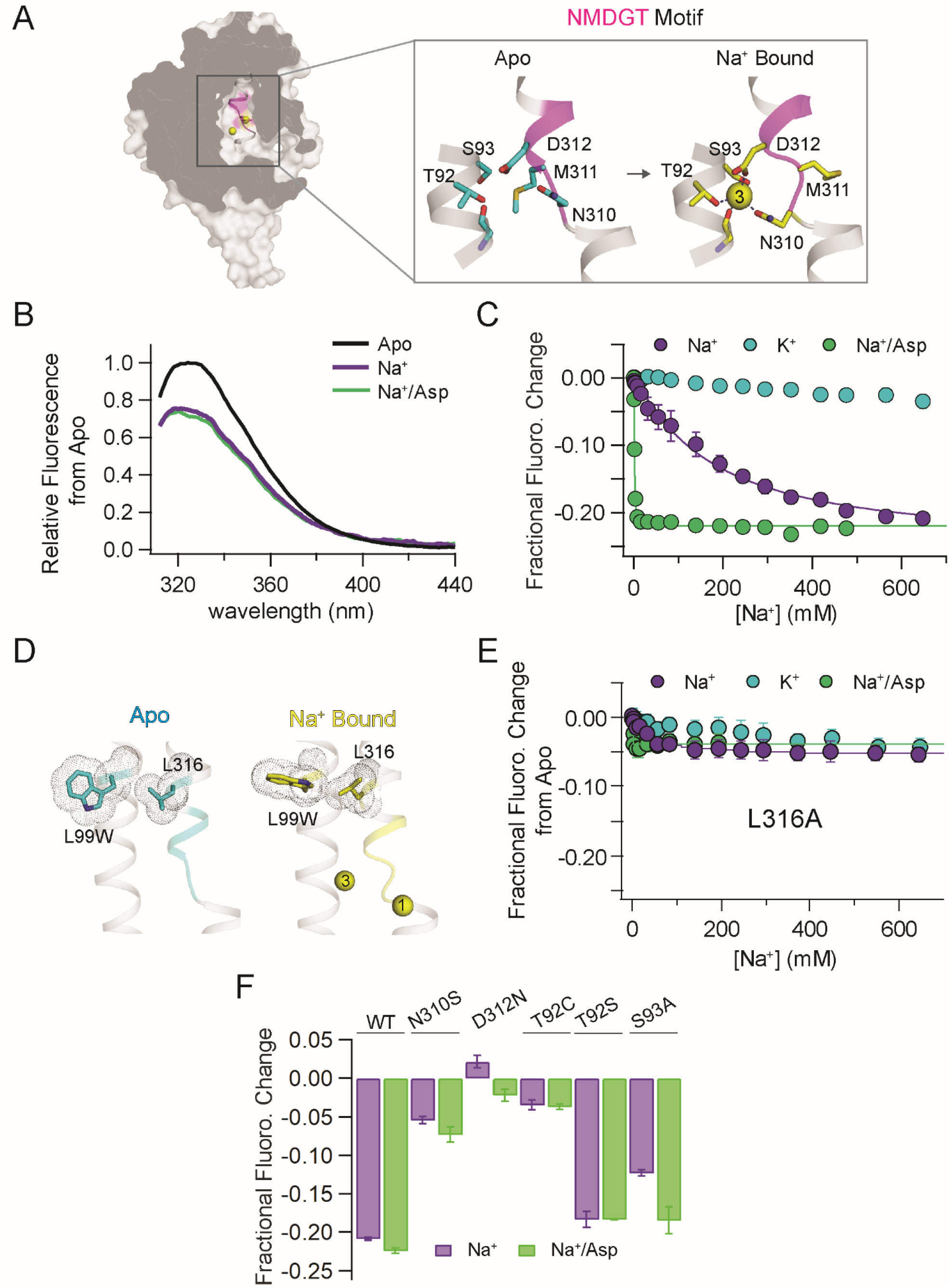
A fluorescence assay for monitoring the NMDGT conformational switch. **A**) Conformational switch in the NMDGT motif on Na^+^ binding. A single protomer of Glt_Ph_ shown in surface representation with the NMDGT motif highlighted in pink. Inset shows the Apo (pdb: 5wdy) and the Na^+^ bound conformations (pdb: 7ahk) of the NMDGT motif. The conformational switch in the NMDGT motif on Na^+^ binding is highlighted by the change in positioning of the M311 side chain. **B**) Emission spectra of Glt_Ph_-L99W on excitation at 295 nm in the following conditions: Apo, 500 mM NaCl and 500 mM NaCl/100 µM Asp. **C**) Titration of L99W with Na^+^, K^+^ and Na^+^ with 100 µM Asp present. The fractional fluorescence change at 324 nm following excitation at 295 nm is plotted and solid lines represent a fit to the Hill equation. **D**) Changes in the positioning of L99W and L316 with a conformational switch in the NMDGT motif. A close-up view of the NMDGT motif in the Apo structure and with Na1 and Na3 bound are shown with L99 mutated to Trp. The W99 and L316 sidechains are shown in space fill representation. Na^+^ ions in A and D are represented as yellow spheres. **E**) Titration of L99W+L316A-Glt_Ph_ with Na^+^, K^+^ and Na^+^ with 100 µM Asp present as in C. **F**) The maximal fluorescence change observed for various Na3 site mutants in the L99W background on titration with Na^+^ and with Na^+^ in the presence of 100 µM Asp. Error bars in panels C, E, F correspond to S.E.M for n ≥ 3.

The structures of Glt_Ph_ in the Apo and Na^+^ bound states indicate that the change in the environment around L99 with the NMDGT conformational switch is due a change in the relative positioning of the L316 side chain (Figure 4D). To test, we investigated the fluorescence changes in a L99W/L316A mutant. We observed that the fluorescence changes were greatly attenuated compared to the L99W mutant (Figure 4E). In control experiments, we confirmed that the L316A substitution did not affect HP2 movement (Figure S3). These experiments confirm that the fluorescence changes of L99W are indeed reporting on the NMDGT conformational switch.

We introduced various Na3 site substitutions into the L99W-Glt_Ph_ and determined the effect on the NMDGT conformational switch (Figure 4F, S4, Table S2). For residues N310 and D312 in the NMDGT motif, we found that the fluorescence changes in the N310S mutant were greatly attenuated compared to the control, L99W-Glt_Ph_ while the D312N substitution eliminated any fluorescence changes with Na^+^ or Na^+^/Asp addition. For the substitutions in T92 in the TM3 helix, we observed substantially reduced fluorescence changes in the T92C mutant while a similar extent of change to the wild type control was observed for the T92S substitution. Consistent with the improved Na^+^ binding observed for the T92S mutant in the Tyr assay, the changes in the NMDGT motif also took place at a lower Na^+^ concentration compared to the control. For the S93A mutant, we observed that the extent of change was similar to the wild type control though the changes were observed at a much higher Na^+^ concentration. We find that substitutions that perturb Na^+^ binding to the Na3 site also effect the NMDGT conformational switch thereby indicating a requirement of Na^+^ binding to the Na3 site for the conformation switch. Further, we find that the effects of the Na3 mutants on the NMDGT conformational switch mirror the effects on HP2 opening, which suggests that the NMDGT conformational switch couples Na^+^ binding at the Na3 site and HP2 opening.

### A S93-M311 steric clash assists in propping HP2 open

M311 in the NMDGT motif is a key residue for coupling Na^+^ and Asp binding in Glt_Ph_.(13, 21) We have previously shown that on Na^+^ binding, M311 acts as a wedge to keep HP2 propped open.(19) In Glt_Ph_ structures, the M311 side chain has been visualized in two conformations (Figure 5A) that we refer to as the extended and the flipped conformation. In the Apo state, the M311 side chain is dynamic and samples both these conformations while in the Na^+^ or the Na^+^/Asp bound state, the M311 side chain is in the flipped conformation. A structural inspection suggests that Na^+^ binding at the Na3 site can influence the conformation of the M311 side chain. This occurs because in the Na^+^ bound conformation, the S93 side chain sterically clashes with the extended conformation of the M311 side chain. To confirm this steric clash mechanism, we probed the effect of substitutions at S93 and M311 that changed the steric properties of these side chains. We substituted S93 with T to increase the steric bulk of this side chain. For the S93T mutant, we observed that the opening of HP2 took place at a lower Na^+^ concentrations and to a larger extent than the wild type (Figure 5B). At M311, a Leu substitution, which has a similar side chain volume (27) to Met but with different sterics, showed HP2 opening in a relatively WT-like manner (Figure 5B). In the M311L/S93T double mutant, we observed that the opening of HP2 took place at a substantially lower Na^+^ concentration and the extent of opening was greater than observed for the single mutants (Figure 5B). These results are in accordance with a steric interaction between the S93 and the M311 side chains when Na^+^ binds to the Na3 site. This steric interaction helps switch the M311 side chain into the flipped conformation, wherein it acts as a wedge to keep HP2 open on Na^+^ binding.

**Figure 5:**
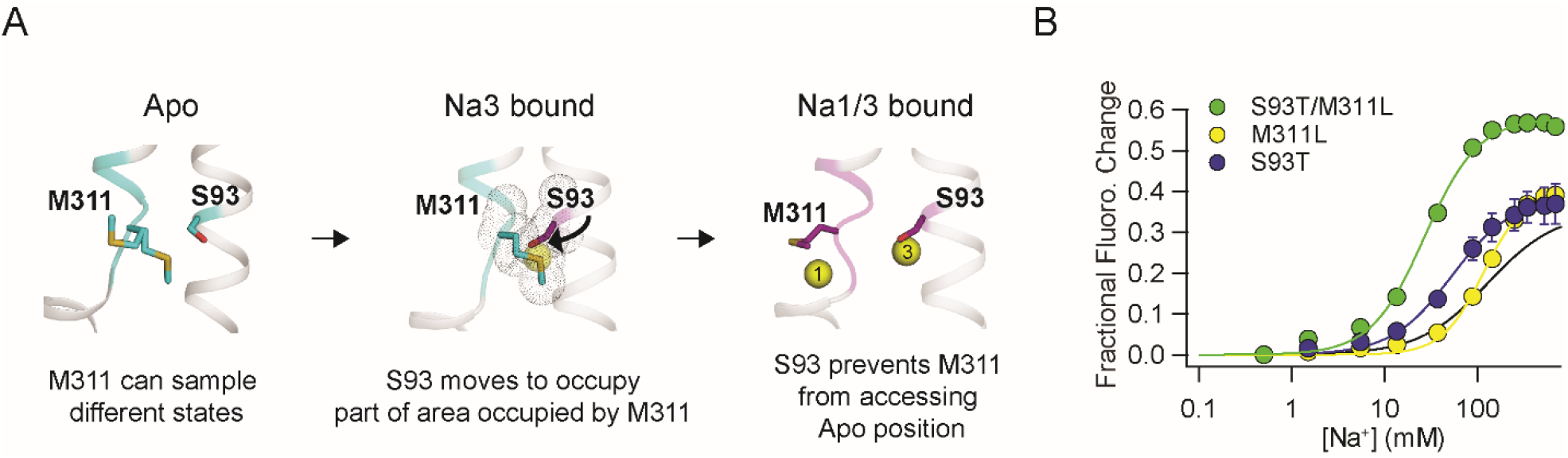
A S93-M311 steric clash at the Na3 site. **A**) Close up view showing the M311 and the S93 side chains in the Apo (left, pdb: 5dwy) and the Na^+^ bound states (right, pdb: 7ahk). Center depicts a hypothetical structure showing the steric clash between the Na^+^ bound conformation of the S93 and the Apo conformation of the M311 with the S93 and M311 side chains shown in van der Waal’s representation. **B**) HP2 opening by Na^+^ in the S93 and M311 mutants. Steady-state titration of the S93T, M311L and the S93T/M311L mutants in the (Phe_CN_+W) background with Na^+^ is plotted and fit (solid line) to give 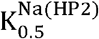 of 53 ± 3.4 mM for S93T, 115 ± 4.7 mM for M311L and 27 ± 2 mM for M93T+M311L. Data for the wild type control are shown as a solid black line. Error bars are S.E.M. for n ≥ 3.

### The D405N substitution at the Na1 site uncouples HP2 opening from Na^+^ binding

We used the D405N substitution to evaluate the effect of perturbing the Na1 site on HP2 movement (Figure 6A). In the HP2 assay, we did not observe a fluorescence change upon the addition of Na^+^ for the D405N mutant although a decrease in fluorescence was observed with Na^+^/Asp (Figure 6B). The decrease in fluorescence indicates Asp binding and the closure of HP2. The lack of a fluorescence signal corresponding to the opening of HP2 with Na^+^ could stem from HP2 not being stable in the open state (as previously observed for the M311A mutant) (19) or due to HP2 already being open. We used TBOA binding to distinguish between these possibilities (as previously described). We found that the addition of Na^+^/TBOA did not result in a significant fluorescence change from the Apo- or the Na^+^-D405N Glt_Ph_ (Figure 6B). For the wild type control, we observed an additional 9% change in intensity when comparing the fluorescence response to Na^+^ versus Na^+^/TBOA. We confirmed that TBOA binds to the D405N-Glt_Ph_ in the presence of Na^+^ by a shift in the Asp affinity in the presence of TBOA (Figure S5). The lack of an effect of Na^+^ or Na^+^/TBOA in the HP2 assay for the D405N-Glt_Ph_ suggests that HP2 is already open in the Apo state. We also investigated a D405A substitution and observed a similar result (Figure S6). We speculated that if HP2 is already open in D405N-Glt_Ph_, then the mutant should be able to bind Asp in the Apo state. We tested Asp binding by the Apo-D405N transporter and observed a decrease in fluorescence indicating that Asp binding and closure of HP2 took place in the absence of Na^+^ (Figure 6B). Asp binding to the Apo D405N transporter took place with a 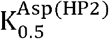 of 124 µM compared to a value of 0.033 µM in the presence of 200 mM Na^+^ (Figure 6C). As anticipated, we did not detect Asp binding to the wild type control in the absence of Na^+^.

**Figure 6:**
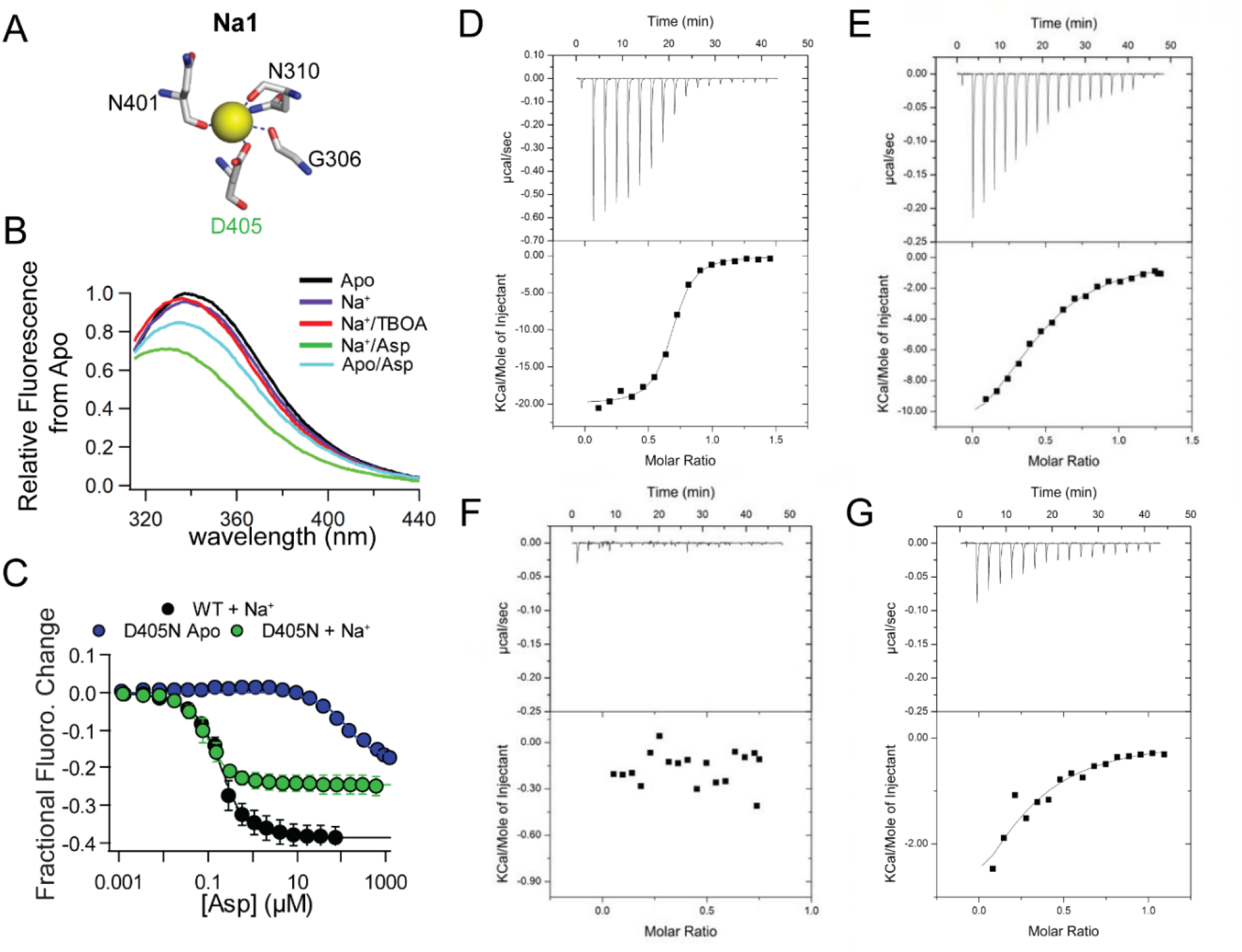
HP2 opening and Asp binding in D405N-Glt_Ph_ is uncoupled from Na^+^ binding. **A**) Close-up view of the Na1 site. **B**) Emission spectra of D405N (Phe_CN_+W) on excitation at 295 nm in the following conditions: Apo, 200 mM NaCl, 200 mM NaCl/10 µM TBOA, 200 mM NaCl/100 µM Asp and Apo/100 µM Asp. **C**) HP2 closure titration with Asp for WT (Phe_CN_+W) in 200 mM NaCl and D405N (Phe_CN_+W) in the Apo state and in 200 mM NaCl. The titration data were fit to give 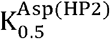 values of 56 ± 2 nM for WT in 200 mM NaCl, 124 ± 21 µM for Apo-D405N and 33 ± 2 nM for D405N in 200 mM NaCl. **D-G**) Asp binding measured using ITC for WT-Glt_Ph_ in 10 mM NaCl (D), D405N in 10 mM NaCl (E), Apo WT (F) and Apo-D405N (G). The upper panels show the heat added to the cell over time with successive additions of Asp and the lower panel shows the integrated heat from each injection plotted as a function of the molar ratio of Asp and Glt_Ph_ in the ITC cell. The solid line represents the fit to the data using an independent binding site model. K_D_ values measured for Asp binding were 0.19 µM for WT-Glt_Ph_ in 10 mM Na^+^, 3.4 µM for D405N in 10 mM Na^+^ and 11 µM for Apo-D405N.

We also used Isothermal Titration Calorimetry (ITC) to evaluate Asp binding to the D405N -Glt_Ph_. In the presence of 10 mM Na^+^, we observed strong Asp binding with a K_D_ of 0.19 µM to the wild type transporter while the Asp binding to the D405N transporter took place with a K_D_ of 3.4 µM and a smaller heat change (Figure 6D/E). We tested Asp binding in the absence of Na^+^ and observed no binding to the Apo wild type control (Figure 6F). In contrast, we observed Asp binding to the Apo D405N transporter with a K_D_ of 11 µM. The magnitude of the heat change on Asp binding to the Apo D405N transporter was smaller than the change observed for Asp binding in the presence of Na^+^ (Figure 6G). The ITC studies confirm that D405N-Glt_Ph_ can indeed bind Asp without Na^+^. However, the lower magnitudes of the heat changes and the shifted affinities indicate differences in the conformational change upon Asp binding in the WT- and the D405N-Glt_Ph_. The combined data from the HP2 assays and the ITC experiments suggest that D405N substitution perturbs the Na^+^ coupled opening of HP2 and allows the binding of Asp to the Apo transporter.

### CW-EPR experiments confirm an open HP2 in D405N-Glt_Ph_

We used CW-EPR experiments to further probe the movements of HP2 when the Na1 site was perturbed. We introduced spin probes at V355 and S279 as previously described to monitor the distance distributions of HP2 in the different states (Figure 7A).(28) The positions at which the spin probes are introduced are the same positions at which fluorescence probes were introduced for the HP2 movement assay. The separation between the spin probes is reflected by the relative amplitude of the central peak in the EPR spectra and in the shape of the spectra.(29) When the spin labels are in close proximity, there is a broadening of the EPR spectra by dipolar relaxation. In wild type Glt_Ph_, HP2 is closed in the Apo state and open when Na^+^ is bound. Correspondingly, we observed an increase in the relative amplitude of the central peak in the EPR spectra of the Na^+^ bound state compared to the Apo state (Figure 7B, C). An additional increase is seen in the Na^+^/TBOA bound state while a decrease is observed in the Na^+^/Asp bound state, which is as anticipated based on the distance distribution of HP2 in these states in wild type Glt_Ph_ (Figure 7C). For the D405N-Glt_Ph_, we observed a similar amplitude for the central peak in the Apo, Na^+^ bound and the Na^+^/TBOA bound states indicating a similar distance distribution for HP2 in these states (Figure 7B, C). We observed a decrease in the relative peak amplitude in the Na^+^/Asp bound state compared to the Apo state, indicating the closure of HP2 on the binding of Na^+^/Asp. However, the change in amplitude between the Na^+^ bound and the Na^+^/Asp state for the D405N transporter was smaller than the corresponding change in the wild type transporter suggesting a limited opening of HP2 in the D405N mutant with Na^+^ compared to the wild type. The splitting pattern in the EPR spectra for the Apo and Na^+^ bound wild type Glt_Ph_ are different, reflecting the different states of HP2. In contrast, we observed a similar splitting pattern for the Apo and Na^+^ bound Glt_Ph_ D405N transporter indicating no change in the state of HP2 in the D405N transporter on the binding of Na^+^. The results from the EPR studies therefore indicate that opening of HP2 in the D405N mutant can take place in the absence of Na^+^ and confirm the findings from the HP2 assays.

**Figure 7:**
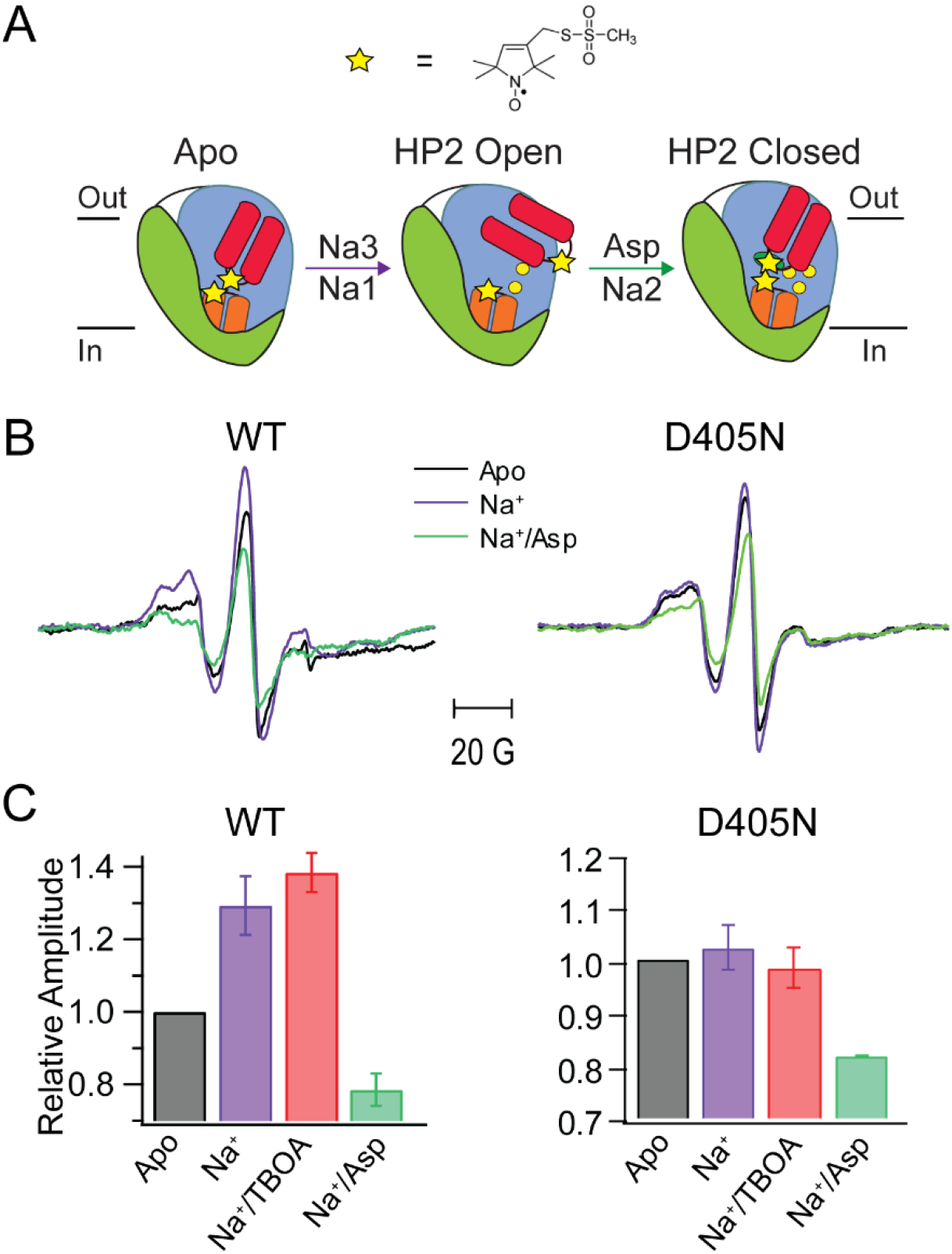
HP2 movement in Glt_Ph_ probed by EPR. A) Cartoon representation of HP2 (red) movement during the coupled binding of Na^+^ ions (yellow spheres) and Asp (green oval). Sites for introducing the MTSL spin probes (yellow stars) for monitoring HP2 movement are 279 in HP1 (orange) and 355 in HP2. **B)** Continuous wave EPR spectra of WT- and D405N –Glt_Ph_ under the following conditions: Apo, 200 mM NaCl, 200 mM NaCl/500 µM Asp. **C**) Relative amplitude for the central peak in the EPR spectra under different conditions normalized to the Apo value. Error bars are S.E.M for n ≥ 3.

### The D405N substitution partially mimics a Na1 only bound state

We have established that HP2 opening in the wild type Glt_Ph_ on Na^+^ binding involves a conformational switch of the NMDGT motif. Next, we probed whether changes in the NMDGT motif are involved in the HP2 opening seen in the Apo-D405N mutant transporter. We were unable to use the L99W substitution for this purpose as the L99W/D405N mutant showed poor biochemistry. A key aspect of the NMDGT switch is a flip of the M311 side chain and in this flipped conformation, the M311 side chain acts as a wedge to keep HP2 propped open on Na^+^ binding. To infer involvement of the NMDGT conformational switch, we evaluated whether the M311 residue plays a role in HP2 opening in the Apo-D405N transporter. We investigated the effect of the M311A substitution in the D405N background. In the M311A/D405N Glt_Ph_(Phe_CN_+W), there is no observable signal for HP2 opening as in the D405N mutant. We inferred the state of HP2 by investigating Asp binding since Asp binding can only take place if HP2 is open. We observed Asp binding to M311A/D405N mutant in the Apo state and the binding took place with a higher affinity than observed for the D405N mutant (Figure 8A). Asp binding to the M311A mutant in the Apo state was not detected.(19) Asp binding to the M311A/D405N mutant in the Apo state indicates that the M311 side chain is not necessary for HP2 opening in the Apo-D405N transporter. This result indicates that the changes in the NMDGT motif and therefore the conformational changes corresponding to Na^+^ binding to the Na3 site are not involved in the HP2 opening seen in the Apo-D405N transporter. In the presence of Na^+^, we observe that the D405N+M311A Glt_Ph_ shows a ∼ 100-fold lower Asp binding affinity compared to D405N Glt_Ph_ (Figure 8B). The change in the affinity with the M311A substitution indicate the participation of the M311 side chain and thereby the NMDGT conformational switch in the Na^+^ coupled binding of Asp to the D405N-Glt_Ph_.

**Figure 8:**
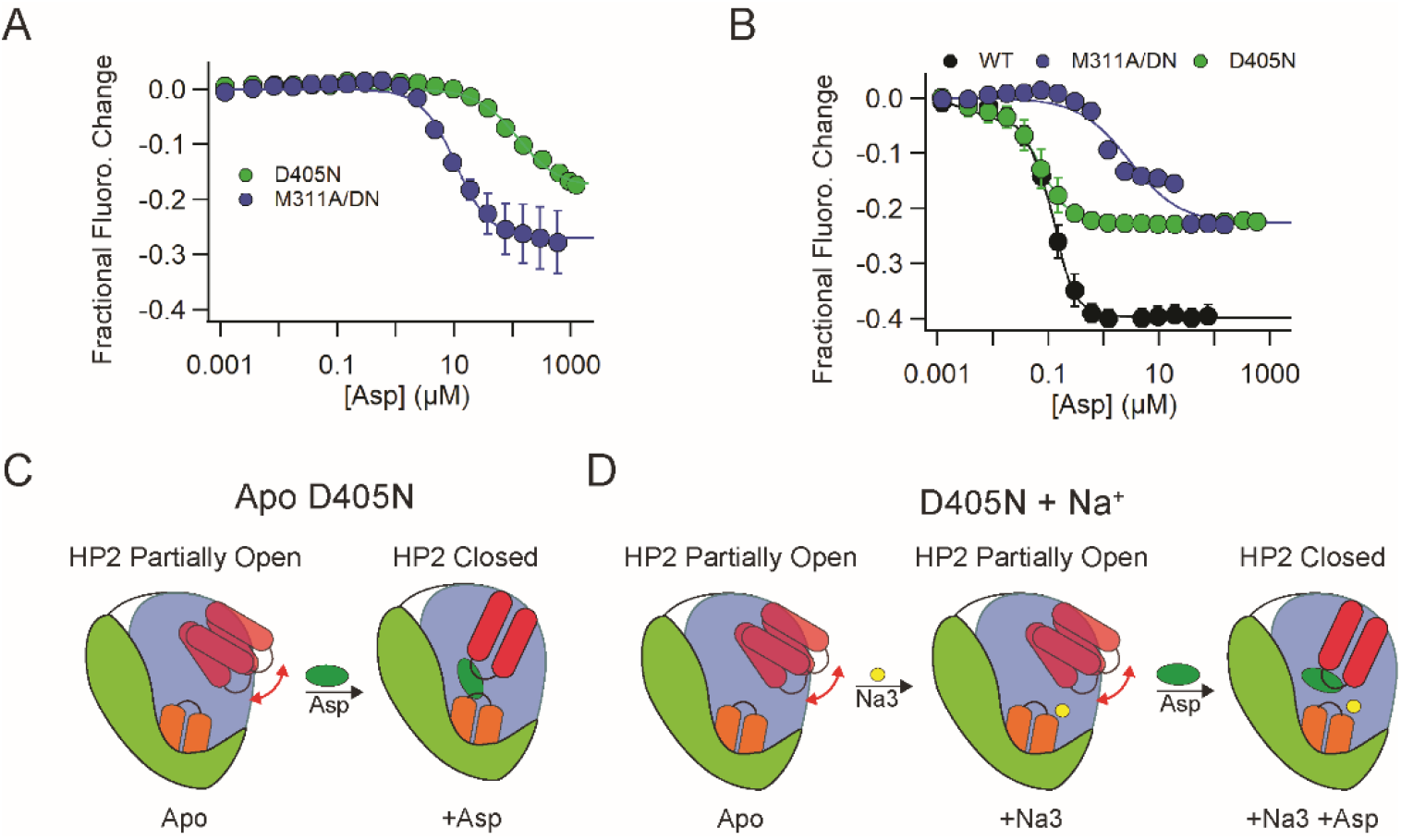
The D405N substitution mimics Na^+^ binding to the Na1 site. **A**) Asp binding to M311A/D405 in the Apo state. Titration of M311A/D405N (Phe_CN_+W) by Asp in the Apo state. The titration data were fit as described in Experimental Procedures to give a 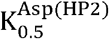 of 11 ± 0.8 µM for M311A/D405N. Titration data for D405N in the Apo state is from Figure 6C. **B**) Titration of WT, D405N and D405N/M311 in the (Phe_CN_+W) background with Asp in 500 mM NaCl. The titration data were fit to give 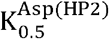 values of 0.017 ± 0.003 µM for WT, 0.021± 0.004 µM for D405N and 2.9 ± 0.9 µM for M311A/D405N. Error bars shown are ± SEM, n ≥ 3. **C**) Cartoon depiction of Asp binding to D405N Glt_Ph_ in the Apo state and in the presence of Na^+^. HP2 in the Apo D405N Glt_Ph_ is in the partially open state and Asp can bind in the absence of Na^+^. Asp binds to the Apo-D405N Glt_Ph_ in a low affinity mode. HP2 is partially open in the D405N mutant and Na^+^ binding to the Na3 site does not increase HP2 opening. Asp binding to the D405N Glt_Ph_ in Na^+^ takes place in the high affinity mode.

The opening of HP2 in the wild type Glt_Ph_ requires Na^+^. HP2 opening in the D405N transporter in the absence of Na^+^ indicates that the D405N substitution partially mimics the effect of Na^+^ binding (Figure 8C). Our experiments with the M311A/D405N mutant have ruled out a participation of the conformational changes corresponding to Na^+^ binding to the Na3 site. As D405 contributes to the Na1 site, we expect that the Asn substitution partially mimics the effects of a Na^+^ ion bound at the Na1 site. Therefore, we conclude that the Apo-D405N Glt_Ph_ resembles a transporter with Na^+^ bound to only the Na1 site. Binding of Asp by the Apo-D405N Glt_Ph_ suggests that HP2 opening and Asp binding can take place with Na^+^ binding to only the Na1 site. This binding of Asp however, takes place in a low affinity mode that is likely distinct from the Asp binding mode seen in the structures of Glt_Ph_ and Glt_Tk_ (Figure 8C). Further, the HP2 assays and the EPR studies indicate a limited HP2 opening when Na^+^ is bound to only the Na1 site in contrast to the full opening of HP2 that is observed when Na^+^ is bound to both the Na1 and the Na3 sites.

## Discussion

The coupled binding of Na^+^ and Asp is central to the transport mechanism in Glt_Ph_. In this study, we developed assays to probe Na^+^ binding to the Na1 and Na3 sites in Glt_Ph_ and to monitor the conformational switch in the NMDGT motif. We used these assays along with the previously developed HP2 movement assay to show that Na^+^ binding to the Na3 site is required for the NMDGT conformational switch while Na^+^ binding to the Na1 site is responsible for the partial opening of HP2. Complete opening of HP2 requires the conformational switch of the NMDGT motif and therefore Na^+^ binding to both the Na1 and the Na3 sites.

Based on our results we propose the following mechanism for HP2 opening by Na^+^ (Figure 9). In the Apo state HP2 is closed. The process of HP2 opening is initiated by Na^+^ binding to the Na1 site. It has been previously suggested that Na^+^ initially binds to the Na3 site as the Na3 site is predicted to have the highest Na^+^ binding affinity.(13, 30, 31) However, the Na3 site is buried within the protein core and Na^+^ access to the Na3 site involves passage through the Na1 site. Computational studies have also previously suggested Na^+^ binding initially to the Na1 site.(31) Occupation of the Na1 site by Na^+^ results in a partial opening of HP2. We anticipate that the partially open conformation of HP2 resembles the HP2 conformation seen in the structure of Glt_Ph_ R397A with Na^+^ present (pdb: 4OYF).(21) The partial opening of HP2 now provides a pathway for Na^+^ ions to access the Na3 site as suggested.(20) Binding of Na^+^ to the Na3 site can take place through this pathway or alternately, the Na^+^ ion at the Na1 site can transition to the Na3 site while the Na1 site is re-occupied by another Na^+^ ion.(31) Occupation of the Na3 site results in a conformational switch of the NMDGT motif which is required for full opening of HP2.

**Figure 9:**
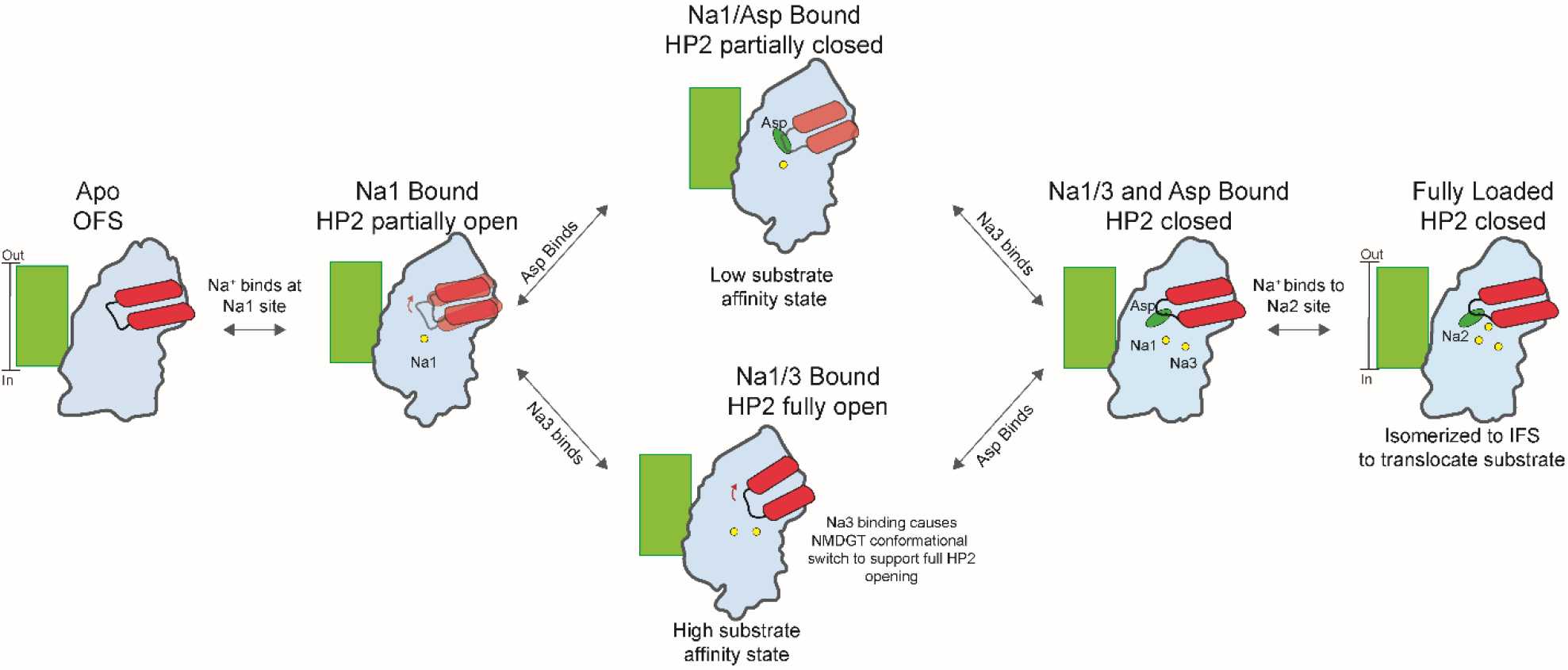
Proposed mechanism for coupled binding of Na^+^ and Asp to Glt_Ph_. HP2 is closed in the Apo state. Na^+^ binding to the Na1 site disrupts some of the interactions that maintain HP2 closed in the Apo state resulting in a partial opening of HP2. Na^+^ then binds to the Na3 site for full opening of HP2 and binding site rearrangements necessary for high affinity Asp binding. Asp binding causes closure of HP2 followed by the binding of Na^+^ to the Na2 site for a fully loaded transport domain. In the alternate pathway, Asp can bind at low affinity to the transporter with Na1 bound. This is followed by Na^+^ binding to the Na3 site which shift Asp binding to the high affinity binding mode and followed by the binding of Na^+^ to the Na2 site resulting in a fully loaded transport domain. The fully loaded transport domain undergoes an elevator like movement for transport and release of Asp and Na^+^ into the cell. Trimerization domain of a protomer is represented as a green box while the transport domain is represented by the blue colored shape. HP2 is shown in red, Asp as the green sphere, and sodium ions as yellow spheres.

Our data suggests that the binding of Asp to Glt_Ph_ can take place through two different pathways. A recent study of a mutant D40N/S279E and the wild GltPh transporter has also reported heterogenous Asp binding consistent with two binding modes for Asp.(32) The conventional pathway consists of Na^+^ binding to the Na1 and the Na3 sites prior to Asp binding. In the alternate pathway, the binding of Asp takes place after the initial binding of Na^+^ to the Na1 site. The binding of Asp is followed by Na^+^ binding to the Na3 site and then to the Na2 site. In the alternate pathway, we anticipate that initial binding mode of Asp with only Na1 bound is a low affinity binding mode that transitions to a high affinity binding mode when Na^+^ ions bind to the Na3 site. We expect that this alternate pathway (Na1→Asp→Na3→Na2) pathway dominates at low Na^+^ and high Asp concentrations, while the conventional pathway (Na3→Na1→Asp→Na2) dominates at high Na^+^ and low Asp concentrations, the physiological conditions under which Glt_Ph_ operates. The presence of this alternate pathway explains how the transporter can bind Asp at low Na^+^ concentrations. A question to be addressed in future studies is visualizing the alternate mode of Asp binding to Glt_Ph_ that dominates at low Na^+^ concentrations. We anticipate that structural studies of the Apo D405N-Glt_Ph_ transporter with Asp bound should reveal this alternate binding mode. Further, we anticipate that the roles Na^+^ binding to the Na1 and the Na3 site on HP2 movement and the pathways for Asp binding elucidated for Glt_Ph_ in this investigation are also operative in EAATs.

## Experimental Methods

### Definition of Terminology used

The Na^+^ dissociation constant determined using the tyrosine fluorescence assay is denoted by 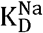. Binding parameters for Na^+^determined using the NMDGT assay are denoted by 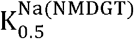 while Na^+^ and Asp binding parameters determined using the HP2 assay are denoted by 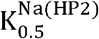 and 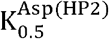 respectively.

### Expression and purification of Glt_Ph_

The Glt_Ph_ construct referred to as wild type consists of seven His substitutions, a C321A substitution and a His_8_ tag at the C-terminus.(9) Amino acid substitutions were introduced by site directed mutagenesis carried out using the PCR overlap or Quickchange mutagenesis protocols. The Glt_Ph_ constructs were cloned and expressed from the pBCH/G4 vector (kindly provided by Dr. Eric Gouaux) in *Escherichia coli* TOP10 cells (Fisher Scientific).(9) Protein expression, membrane preparation and protein purification using affinity chromatography and size exclusion chromatography (SEC) was carried out as previously described.(33) SEC was carried out using a Superdex S-200 column (Cytiva) using 15 mM Tris-HEPES pH 7.5, 100 mM NaCl, 1 mM EDTA and 0.1% (w/v) DDM (Na^+^ -SEC buffer) as the column buffer. For Glt_Ph_ proteins used for the Na^+^ assays, the Na^+^ present was removed after affinity chromatography by concentration using an Amicon filter (100 kDa cutoff) and extensive washing with 20 mM Tris-HEPES pH 7.5, 100 mM KCl, 1 mM EDTA and 0.1% (w/v) DDM (K^+^- SEC buffer) to ensure that the Na^+^ concentration was below 1 mM. The Glt_Ph_ proteins were then further purified by SEC using the K^+^-SEC buffer.

### Unnatural amino acid incorporation

Incorporation of Phe_CN_ (4-Cyano-Phenylalanine) was accomplished using the PheCN tRNA synthetase/tRNA pair on a pULTRA plasmid as previously described with the exception that 1 mM of PheCN (Chem-Impex) was used in the expression media.(19, 34) The pULTRA-CNF plasmid was a gift from Dr. Peter Schultz, Addgene plasmid # 48215. The spectinomycin resistance gene on the pULTRA-CNF plasmid was replaced by the kanamycin resistance gene from pRSF1b (EMD Millipore) for use in TOP10 cells.

### Tyrosine-based Na^+^ binding assay

Steady state Na^+^ titrations were carried out with ∼40 µgs of Glt_Ph_ in 20 mM Tris-HEPES pH 7.5, 200 mM Choline chloride, and 0.1% DDM (choline buffer) at 30 °C. The change in fluorescence on Na^+^ addition was monitored at 308 nm after excitation at 289 nm following an incubation time of 1 min. The fluorescence changes measured were corrected for dilution and the fractional fluorescence change from the Apo protein (F) was fit to the Hill equation (Equation 1).

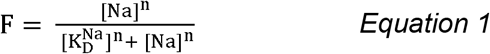

### NMDGT conformational switch assay

Steady-state NMDGT assays were carried out with ∼80 µgs of Glt_Ph_ in choline buffer at 30 °C. The change in fluorescence with Na^+^ addition was monitored at 324 nm after excitation at 295 nm following a 1 min incubation. The fluorescence change measured was corrected for dilution and the fractional fluorescence change from the Apo protein was fit to a Hill equation (*Equation 1*) for determination of 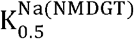.

### Hairpin 2 movement assay

HP2 movement assays were carried out using Glt_Ph_ transporters with a V335W and S279Phe_CN_ along with the desired amino acid substitution/s in the choline buffer at 30 °C. The changes in fluorescence at 345 nm were monitored after excitation at 295 nm. The HP2 opening titrations were carried out by adding aliquots of Na^+^ to the Apo transporter and monitoring the change in fluorescence after 10 min. incubation. The fluorescence change was corrected for dilution and fit using a Hill equation (equation 1) to determine the value of 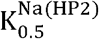. The HP2 closing titrations were carried out by adding Asp to the protein in the presence of Na^+^ and monitoring the change in fluorescence after a 1 min. incubation. The change in fluorescence (F) was corrected for dilution and fit using equation 2 to determine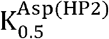.

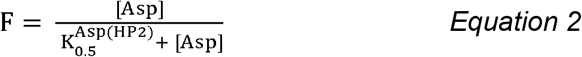

Equation 3 was used in cases where the protein concentration (P) was similar to or greater than the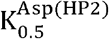.

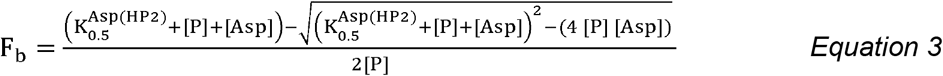

### Dye-based Na^+^ binding assay

Na^+^ binding to Glt_Ph_ constructs was assayed using the voltage sensitive dye, RH421 as previously described.(18, 21) Briefly, 200 nM of RH421 was added to 300 μgs of Glt_Ph_ in 10 mM Tris-HEPES pH 7.5, 100 mM KCl, 0.02% (w/v) DDM. Following dye addition, the protein solution was allowed to equilibrate (∼1 hour) before the start of the titration.

Aliquots of NaCl were added and Na^+^ binding was monitored by the change in the fluorescence emission at 628 nm following excitation at 532 nm. After the addition of each aliquot of NaCl, the sample was allowed to equilibrate (∼ 15 min) until the fluorescence signal stabilized. For Na^+^ titrations in the absence of Asp, the maximal fluorescence change was determined at the end of the titration by the addition of 1 mM Asp. The fluorescence changes measured were corrected for dilution, normalized to maximal fluorescence change observed and fit to a Hill equation (Equation 1) to determine the K_D_.

### Isothermal Titration Calorimetry

The Glt_Ph_ proteins used for the ITC experiments were purified as described. Following affinity chromatography, the His_8_ tag present at the C-terminus was removed by proteolysis with Thrombin and the cleaved proteins were purified by SEC using 20 mM Tris-HEPES pH 7.5, 100 mM KCl, and 0.4 mM (use %) DDM as the column buffer. ITC experiments were carried out with 15-40 µM of the protein in the respective assay buffer (20 mM Tris-HEPES pH 7.5, 100 mM KCl, 0.4 mM DDM or 20 mM Tris-HEPES pH 7.5, 90 mM KCl, 10 mM NaCl, 0.4 mM DDM). The experiments were carried out at 25 °C using an ITC200 calorimeter (Malvern Panalytical). The titrant solution of Asp was made in the corresponding assay buffer. Data was analyzed using MicroCal Origin-based software and an independent-binding site model as previously described.(18, 35).

### EPR Spectroscopy

The double cysteine Glt_Ph_ mutants (V355C/S279C or V355C/S279C/D405N) were expressed and purified as described. Following affinity chromatography, the Glt_Ph_ proteins were reduced with 500 µM tris(2-carboxy-ethyl) phosphine (TCEP, Gold Biotechnology) for 30 min at room temperature. The proteins were then concentrated using an Amicon filter (100 kDa cutoff) and washed extensively with 20 mM Tris-HEPES pH 7.5, 50 mM NaCl, 0.1% DDM for removal of TCEP and imidazole. The protein concentrations were determined and the proteins were labelled with the spin probe (1-oxyl-2,2,5,5-tetramethylpyrrolidin-3-yl) methyl methanethiosulfonate (MTSL, Toronto Research) at a 10: 1 probe-to-protein ratio.(28) The labeling was carried out for 30 min at room temperature followed by overnight incubation at 4 °C. The labeled protein was separated from free label by SEC in K^+^-SEC buffer and a subsequent concentration step with an Amicon filter (100 kDa cutoff). The spin-labelled protein was incubated with the respective ligands (Na^+^, Asp or TBOA) and EPR spectra were collected with ∼100 µM labelled protein on a Bruker E500-X band EPR spectrometer equipped with a super X microwave bridge and an ER4123D dielectric resonator (Bruker Biospin). EPR spectra were collected using the following parameters: microwave frequency of 9.78 GHz, microwave power of 2mW, modulation frequency of 100 kHz, and modulation amplitude of 2G.

EPR spectra were baseline subtracted and double integrated to determine the number of spins present in each sample. Spectra were normalized to the spin number for comparison between the different conditions. Relative amplitudes were determined by subtracting the central peak minima from the maxima.

### Aspartate transport assays

Glt_Ph_ was reconstituted into lipid vesicles comprised of a 3: 1 ratio of *E*.*coli* polar lipids to 1-palmitoyl-2-oleoyl-glycero-3phosphatidylcholine (POPC) as previously described.(36) The transport assays were carried out in 20 mM Tris-HEPES pH 7.5, 200 mM NaCl, 1 µM valinomycin at 30 °C using 100 nM of ^14^C Asp (Moravek Biochemicals) as previously described.(33) Background levels of Asp transport were determined by performing the transport assay in absence of NaCl, with 100 mM KCl in the assay buffer.

## Acknowledgements

This research was supported by NIH Grants R01GM087546, R21NS113561 (to F.I.V) and R37NS085318 (to Dr. Olga Boudker, Principal Investigator and F.I.V., Co-investigator). E.A.R. was supported by a Predoctoral Fellowship from the American Heart Association (AHA 19PRE34380950). We thank Drs Krishna Reddy, Olga Boudker, Lucy Forrest and José Faraldo-Gómez for helpful discussions, Dr. Eric Gouaux for access to the ITC, Dr Vikas Navratna for his generous training in use of ITC, and Dr. Steve Lockless for help with analysis of ITC data.

## Notes

### Competing Interest Statement

The authors have declared no competing interest.

