## Supplementary Materials for "Distinct roles of the Na^+^ binding sites in the allosteric coupling mechanism of the glutamate transporter homolog, Glt_Ph_"

A

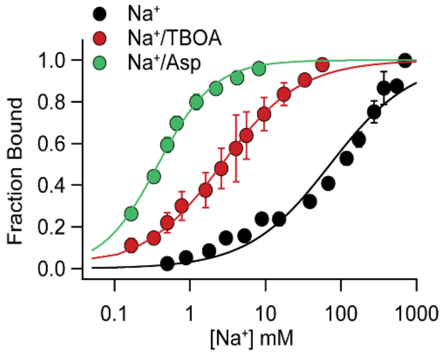


B

| Site | Substitution | $K_D$ (mM) | Fold Change from WT Dye | Fold Change from WT Tyr | $K_D$ (mM) +1 mM Asp |
|---|---|---|---|---|---|
| | WT | 72 ± 10 | | | 0.4 ± 0.02 |
| Na3 | S93A | 352 ± 40 | 4 | ~6 * | 3.9 ± 0.2 |
| Na3 | T92C | 237 ± 24 | 3 | 3 | 2.5 ± 0.2 |
| Na3 | N310S | 279 ± 17 | 3 | 3 | 2.5 ± 0.1 |
| Na3 | D312N | 234 ± 19 | 3 | 2 | 1.3 ± 0.1 |
| Na1 | D405N | 224 ± 28 | 3 | 4 | 1.6± 0.1 |

**Figure S1**. **$Na^+$ affinity of the Na1 and Na3 site mutants using the dye-based assay. A)** Steady state $Na^+$ binding curves under different conditions ($Na^+$, $Na^+$/10 µM TBOA and $Na^+$/1 mM Asp) **B)** Table of $Na^+$ affinities for Na1 and the Na3 site mutants using the dye-based assay. The fold changes in affinity for the Na site mutants determined using the dye-based assay and the tyrosine fluorescence assay are shown. (*) Saturation was not observed for the S93A mutant and the fold change reported is a rough estimation. Errors in A and B are S.E.M. for n ≥ 3.

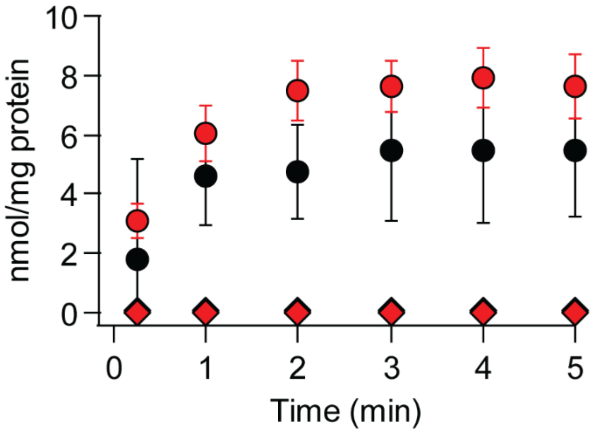


**Figure S2**. **Asp uptake activity by L99W-Glt$_{Ph}$.** The time course of [$^{14}$C] L-Asp uptake by WT-Glt$_{Ph}$ (black circles) and L99W-Glt$_{Ph}$ (red circles) in the presence of a $Na^+$ gradient. No uptake is observed for the WT (black diamonds) or L99W-Glt$_{Ph}$ (red diamonds) in the absence of a $Na^+$ gradient. Error bars for the uptake data points for L99W-Glt$_{Ph}$ in $Na^+$ are S.E.M. for n =3 while the error bars for the other three samples represent the range of values for n= 2.

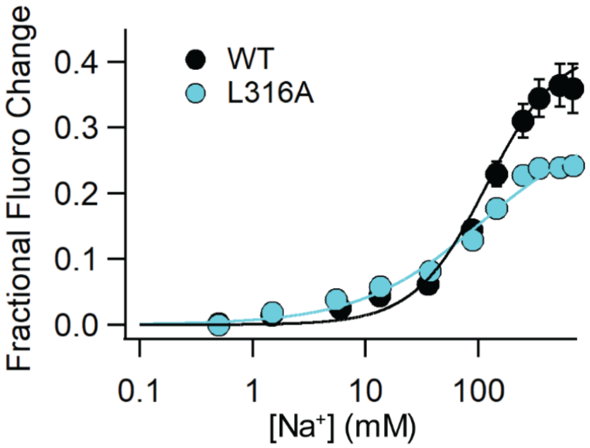


**Figure S3. HP2 movement in L316A-Glt$_{Ph}$.** Steady-state titration of L316A-Glt$_{Ph}$(Phe$_{CN}$+W) and the wild type control by $Na^+$ is plotted and fit (solid line) to give a $K_{0.5}^{Na(HP2)}$ of 106 ± 45 mM for L316A and 132 ± 20 mM for the wild type. Error bars are S.E.M. for n ≥ 3.

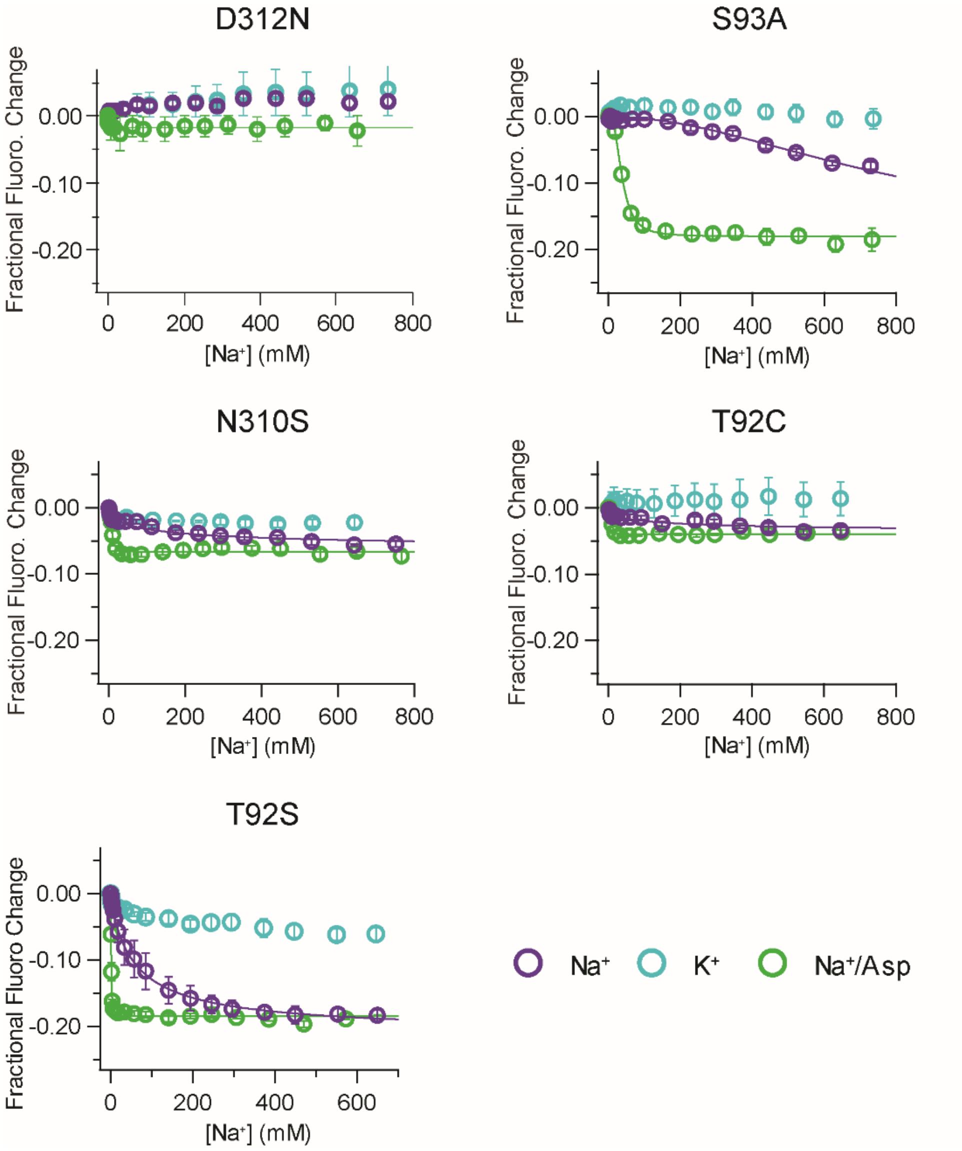


**Figure S4. The NMDGT conformational switch in the Na3 site mutants.** Steady state $Na^+$ titration data for Na3 site mutants using the NMDGT conformational switch assay. Solid lines represent a fit to the Hill equation used to determine the $K_{0.5}^{Na(NMDGT)}$ values.

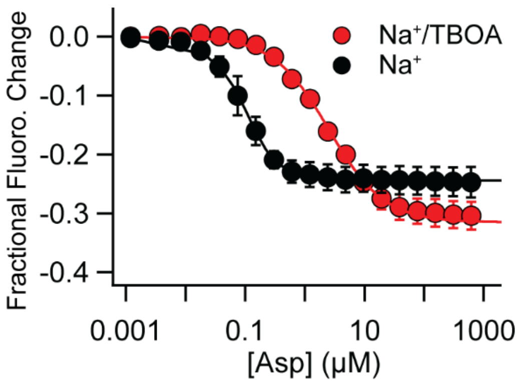


**Figure S5. TBOA binding by D045N-Glt$_{Ph}$.** Steady state titrations of D405N-Glt$_{Ph}$(PheCN+W) in 200 mM $Na^+$ by Asp or by Asp with 100 µM TBOA present gave $K_{0.5}^{Asp(HP2)}$ values of 33 ± 2 nM in the absence of TBOA and 2300 ± 80 nM in the presence of TBOA The ~70 fold change in Asp affinity in the presence of TBOA indicates TBOA binding by the D405N mutant. All error bars are S.E.M. for n ≥ 3.

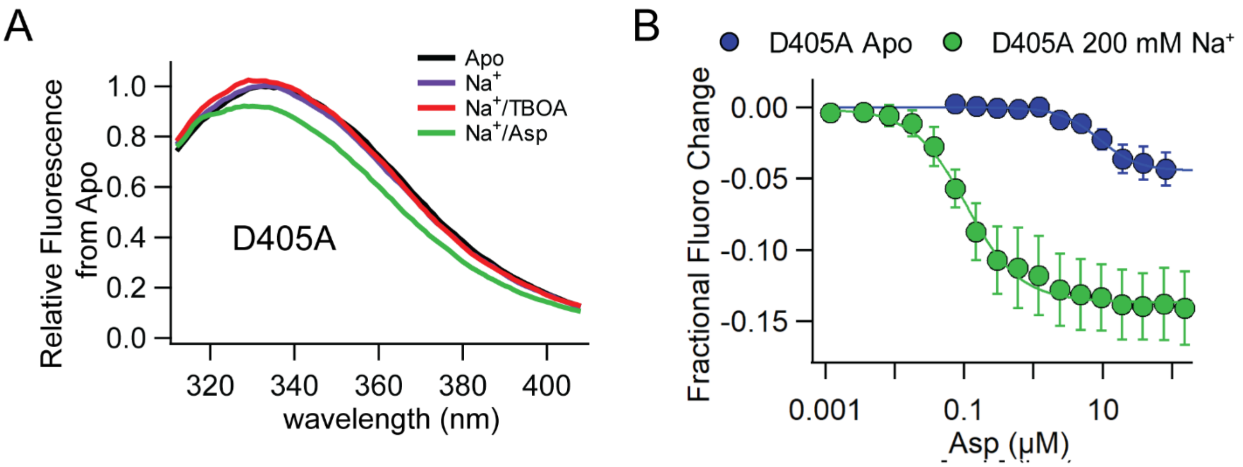


**Figure S6. HP2 opening in D405A-Glt$_{Ph}$.** **A)** Fluorescence emission spectra for D405A-Glt$_{Ph}$(Phe$_{CN}$+W) on excitation at 295 nm in the following conditions: Apo, 200 mM NaCl, 200 mM NaCl/10 µM TBOA and 200 mM NaCl/100 µM Asp. **B)** Steady state titrations of D405A by Asp with and without $Na^+$ (Apo) present using the HP2 movement assay gave $K_{0.5}^{Asp(HP2)}$ values of 84 ± 2 nM for D405A+$Na^+$ and 19.8 ± 6 µM for D405A-Apo. All error bars are S.E.M. for n ≥ 3.

**Table S1**: $Na^+$ binding affinities for the Na site mutants using the tyrosine-fluorescence based assay.

| | | $K_D^{Na}$ (mM) | Hill Coefficient |
|---|---|---|---|
| WT | | 44.7 ± 11 | 1.1 ± 0.2 |
| Na3 | S93A* | > 300 | |
| | S93C* | 304 ± 22 | 2.2 ± 0.2 |
| | T92C | 139 ± 62 | 1 ± 0.4 |
| | T92S | 14 ± 2.8 | 1.3 ± 0.01 |
| | N310S | 171 ± 32 | 1 ± 0.2 |
| | N310D | 178.8 ± 22 | 1.3 ± 0.3 |
| | D312N | 123 ± 11 | 1.1 ± 0.2 |
| Na1 | D405N | 214 ± 18 | 1.7 ± 0.2 |

(*) S93A, C did not saturate and $K_D^{Na}$ values reported are approximate. Errors are S.E.M. and n ≥ 3.

**Table S2.** Apparent $Na^+$ affinities and Hill Coefficients for the Na3 site mutants measured using the NMDGT assay.

| | $Na^+$ | | $Na^+$/ Asp | |
|---|---|---|---|---|
| | $K_{0.5}^{Na(MDGT)}$ (mM) | Hill Coefficient | $K_{0.5}^{Na(NMDGT)}$ (mM) | Hill Coefficient |
| WT | 172 ± 6 | 1.1 ± 0.05 | 2 ± 0.06 | 2.2 ± 0.1 |
| S93A | 804 ± 15 | 2 ± 0.1 | 38 ± 1.1 | 2.6 ± 0.2 |
| T92C | 102 ± 23 | 0.6 ± 0.1 | 8.2 ± 0.4 | 3 ± 0.4 |
| T92S | 60 ± 4.4 | 0.9 ± 0.04 | 1.5 ± 0.06 | 1.8 ± 0.1 |
| N310S | 108 ± 14 | 0.6 ± 0.05 | 7.7 ± 0.6 | 2.4 ± 0.4 |
| D312N | No change | | 1.8 ± 0.7 | 1.3 ± 0.7 |

Errors are S.E.M. for n ≥ 3.
